# Gibberellic acid and light effects on seed germination in the seagrass *Zostera marina*

**DOI:** 10.1101/2024.07.16.603723

**Authors:** Riccardo Pieraccini, Lawrence Whatley, Nico Koedam, Jasper Dierick, Ann Vanreusel, Tobias Dolch, Tom Van der Stocken

## Abstract

- Seagrass meadows have been heavily affected by human activities, with *Zostera marina* being one of the most impacted species. Seed-based methods are currently the preferred approach for their restoration. However, low germination rates and seedling establishment often affect the success rate and feasibility of restoration projects.
- We tested, for the first time, the combined effect of light spectra (white and red light and darkness), photoperiod, and gibberellic acid (GA_3_) on seed germination rates in *Z. marina*, by means of an incubation experiment with a fully crossed design, employing penalised logistic regression and Cox proportional hazards analysis. Seedling development was subsequently monitored to assess the potential side-effects of the priming agents on morphometric growth.
- Light priming positively affects germination, with germination probability being substantially increased when red light and darkness treatments were combined with GA_3_. Time to germination was reduced at mid- to high- GA_3_ concentrations. Morphometric analysis of the cotyledonary and leaf tissue development did not indicate *a posteriori* side-effects of seed priming on growth.
- Light and GA_3_ priming favour germination probability and release from dormancy in *Z. marina* seeds. Seed priming can reduce stress- or manipulation-induced dormancy and can be considered in contexts where on-demand germination is required.

## Introduction

Seagrasses are marine flowering plants widely distributed along the world’s coastal areas (Short *et al*., 2007a). They provide valuable ecosystem services to coastal communities and climate-change mitigation, including carbon sequestration (Serrano *et al*., 2021), nursery and foraging habitat for marine organisms (McDevitt-Irwin *et al*., 2016), supporting global fisheries (Unsworth *et al*., 2019a), as well as coastal protection from erosion through sediment accumulation and stabilisation (Duarte, 2002). Unfortunately, seagrass ecosystems are facing increasing pressure from human activities and climate-change effects, with reported global loss accelerating from 0.9% per year in 1940 to 7% per year since 1990 (Short *et al*., 2007b; Waycott *et al*., 2009; Unsworth *et al*., 2019b). Restoration initiatives are currently being carried out worldwide to rehabilitate these ecosystems and offset the biodiversity loss of the habitat associated with seagrass meadows (Van Katwijk *et al*., 2016; Jacob *et al*., 2018; Valdez *et al*., 2020). Seagrass conservation and restoration efforts are increasingly prioritized by international bodies such as the United Nations (UNEP, 2020) and the European Union (COM, 2024), emphasizing the urgent need to protect and restore these vital marine habitats. These initiatives aim to strengthen restoration efforts and support ecosystem resilience.

*Zostera marina*, the most wide-ranging seagrass species, has been the subject of numerous restoration attempts in the last decades (Infantes *et al*., 2016; Yang *et al*., 2016). Seed-based methods are currently among the favoured restoration techniques (Unsworth *et al*., 2019), as they offer several advantages, including minimal disruption to donor meadows (Zhang *et al*., 2022), increased genetic diversity (Sinclair *et al*., 2013), and reduced logistic constraints, as observed for wild-sourced sod transplantation (Unsworth *et al*., 2023).

Seed broadcasting holds the potential to facilitate a rapid expansion of seagrass meadows, with successful projects reporting very high growth rates and increases of over 16 times the original meadow extent (from 213 ha to 3612 ha of newly vegetated habitat) (Orth *et al*., 2020). Additionally, new advancements in seed broadcasting techniques have significantly enhanced the effectiveness and scalability of seagrass restoration efforts over the past decade, including seed injection (Gräfnings *et al*., 2023), seagrass seed line (Unsworth *et al*., 2019a), and rewilding approaches (Van Katwijk *et al*., 2016).

The multiplication of restoration projects aimed at counteracting the degradation of *Z. marina* meadows has resulted in an exponential increase in seed demand (Unsworth *et al*., 2023). To support this high demand, many new *Z. marina* nurseries have been established worldwide; however, seed production remains limited (Global Seagrass Nursery Network, personal communication). Thus far, most seagrass restoration efforts rely on wild-collected seeds (Unsworth *et al*., 2023). Therefore, it is essential to optimise and maximise the use of the available seed stock.

Seed production represents a substantial energy investment for seagrass populations, with yields reaching up to 10 million seeds per hectare (Statton *et al*., 2017). These seeds hold high viability upon release, a natural strategy to support meadow regeneration (Riddin & Adams, 2009; Jarvis & Moore, 2010; Reynolds *et al*., 2013). However, studies conducted across the world consistently report poor seagrass regeneration from seed-based restoration, with more than 90% of seeds failing to germinate and/or establish successfully (Marion & Orth, 2010b; Orth *et al*., 2003, 2007). Stresses induced during seed collection, manipulation, and storage for restoration purposes can significantly affect seed batch quality (Orth *et al*., 2000; Turner *et al*., 2013), prolonged dormancy (Orth *et al*., 2000) and leading to reduced germination (Probert *et al*., 2009; Dooley *et al*., 2013).

Seed dormancy is an evolutionary adaptation conferring seed embryos the capacity to endure unfavourable environmental conditions *prior* to germination (Finch-Savage & Leubner-Metzger, 2006; Yan & Chen, 2020), thereby allowing seed maturation and overwintering (Zhu *et al*., 2023). Dormancy modulation is intricately linked with environmental cues, where changes in temperature, salinity, irradiance, and photoperiod are typically associated with seasonal shifts (Batlla & Benech-Arnold, 2010; Footitt *et al*., 2017). Photostimulation is observed in many angiosperm taxa (Koller, 1957; Botha & Small, 1988; Thanos *et al*., 1991, 1994; Carta *et al*., 2017; Vandelook *et al*., 2018; Mérai *et al*., 2019) and is considered to act as a catalyst of germination, directly influencing the physiological activities of dormant seeds with the release from dormancy and the promotion of germination (Finch-Savage & Leubner-Metzger, 2006; Yan & Chen, 2020).

Plants detect and transduce light signals through various photoreceptors (Gyula *et al*., 2003; Li *et al*., 2011), with a recent study revealing that the genome of *Z. marina* still contains the genes associated with six of these photoreceptors (Olsen *et al*., 2016). The activation of photoreceptors initiates a series of physiological processes involved in the regulation of hormonal pathways and hormone biosynthesis (Sharrock, 2008; Seo *et al*., 2009; Panda *et al*., 2022). Abscisic acid (ABA) is considered the principal hormone driving the initiation and maintenance of seed dormancy (Koornneef *et al*., 2002). Gibberellins (GAs), in turn, break dormancy and promote germination. Increased levels of GA within the embryo initiate enzymatic processes that promote the degradation of the endosperm (Kahn *et al*., 1957; Oh *et al*., 2006; Toyomasu *et al*., 1998), thereby supporting embryo development by facilitating the rupture of the testa and the emergence of the cotyledon (Finch-Savage & Leubner-Metzger, 2006). Red- and far-red light receptors, known as phytochromes, appeared to be involved in the germination process of many angiosperms (Panda *et al*., 2022; Seo *et al*., 2009; Sharrock, 2008). Phytochrome responses involve the conversion of its inactive form, Pr, to its bioactive form, Pfr, upon absorption of red light, thereby initiating a signalling cascade (Furuya & Schäfer, 1996).

In some seagrass species, an increase in germination rates is observed upon seed exposure to red- and far-red light, such as in *Halophila ovalis* (Strydom *et al*., 2017; Waite *et al*., 2021), *Thalassia hemprichii* (Soong *et al*., 2013), and partially in *Zostera marina* (Wang *et al*., 2017). However, recent advances in *Z. marina* genomics revealed an alteration of these photo-hormonal pathways, including the absence of some red and far-red receptors, indicating that the response to light spectra may differ in *Z. marina* compared to other plant species (Olsen *et al*., 2016).

In agriculture and terrestrial restoration, it is common practice to expose seeds to stimuli, such as light and hormone baths, designed to release from dormancy and increase germination success (Dutta, 2018; Chua *et al*., 2020; Kettenring & Tarsa, 2020). Similarly, in seagrasses, seeds are sometimes exposed to preliminary treatments conducive to germination, such as seed scarification, temperature, and/or freshwater pulses (Kaldy, 2014; Statton *et al*., 2017; Liu *et al*., 2023; Wang *et al*., 2017); however, due to the complex and variable nature of seeds and their maturation, outcomes often yield mixed results, hindering the establishment of a replicable protocol.

Few studies have directly explored the effect of light priming on the germination of *Z. marina* seeds (Moore *et al*., 1993; Wang *et al*., 2017); however, to the best of our knowledge, no studies have yet considered hormone priming and its combination with light spectra to induce seed germination and release dormancy.

Light and phytohormone priming have the potential not only to enhance germination rates but also to induce the pre-germinative stage without triggering actual sprouting prior to sowing, ensuring quicker and more uniform germination of the seed stock. Therefore, in this study, we investigate seed germination in a North Atlantic population of *Z. marina*, focusing on seeds’ physiological responses to different (1) light spectra, (2) photoperiods, and (3) GA_3_ concentrations. We examine how these factors influence seed germination rates and explore temporal patterns in germination. Our overarching goal is to identify priming treatments that facilitate on-demand germination. Additionally, the morphometric characteristics of seedling development (post-germination) are analysed to assess the potential growth side-effects of the priming agents.

## Material and methods

### Seed collection and storage

Seed bearing shoots of *Zostera marina* were collected directly from the seagrass meadows in Hamburger Hallig (Germany) during low tide in September 2021. A permit was released by Landesbetrieb für Küstenschutz, Nationalpark und Meeresschutz Schleswig-Holstein (Germany), on June 25^th^, 2021 (n. 3141-537.46). Shoots were transported within 28 hours to the lab facility at Ghent University in refrigerated boxes (8 ± 1°C).

Seed release and ripening were ensured in aerated buckets (30 L, 30 psu, 10 ± 0.5°C) for a period of 42 days; then seeds were hand-sorted from detritus and gently rinsed prior to the dormancy period. Winter dormancy (i.e., cold stratification) was simulated by incubating the seeds in full dark conditions at 3 ± 0.5°C for a total of 120 days. Autoclaved natural seawater (121 °C, 15 psi, 20 min) with adapted salinity (28 psu), hereafter referred to as SSW, was used as media and storing solution. SSW was supplemented with 0.2 mg L^-1^ copper sulphate (CuSO_4_) to prevent oomycete infection, as suggested by Govers *et al*. (2017), and the solution was refreshed weekly.

Seeds were subjected to a viability test, using a non-destructive method described by Marion & Orth (2010) (e.g., fall velocity, colour, firmness, etc.). Seeds that were considered viable upon this viability test were surface-sterilised with ethanol (EtOH 70% v/v) for 2 min and subsequently with a solution of 1% sodium hypochlorite (NaOCl) for 5 min. Seeds were then rinsed five times in SSW.

### Experimental design, culture media, and culture condition

A fully crossed experimental design was adopted to assess the effect of light spectra, photoperiod, and hormone priming on the germination of *Z. marina* seeds (Fig. **1**). Five hormone priming solutions (culture media) were prepared by dissolving various concentrations of commercially available GA_3_ (Sigma-Aldrich), a major bioactive form of gibberellic acid. The concentrations tested included 10 mg L^-1^, 50 mg L^-1^, 500 mg L^-1^, and 1000 mg L^-1^ in SSW and a negative control with 0 mg L^-1^ in SSW. Phytohormone solutions were mechanically filtered under laminar flow using sterile 0.2 µm syringe filters (Pall, Acrodisc). Excess solutions were stored at -20 ± 0.5°C and used for the weekly refreshment of the experimental units.

**Fig. 1.**
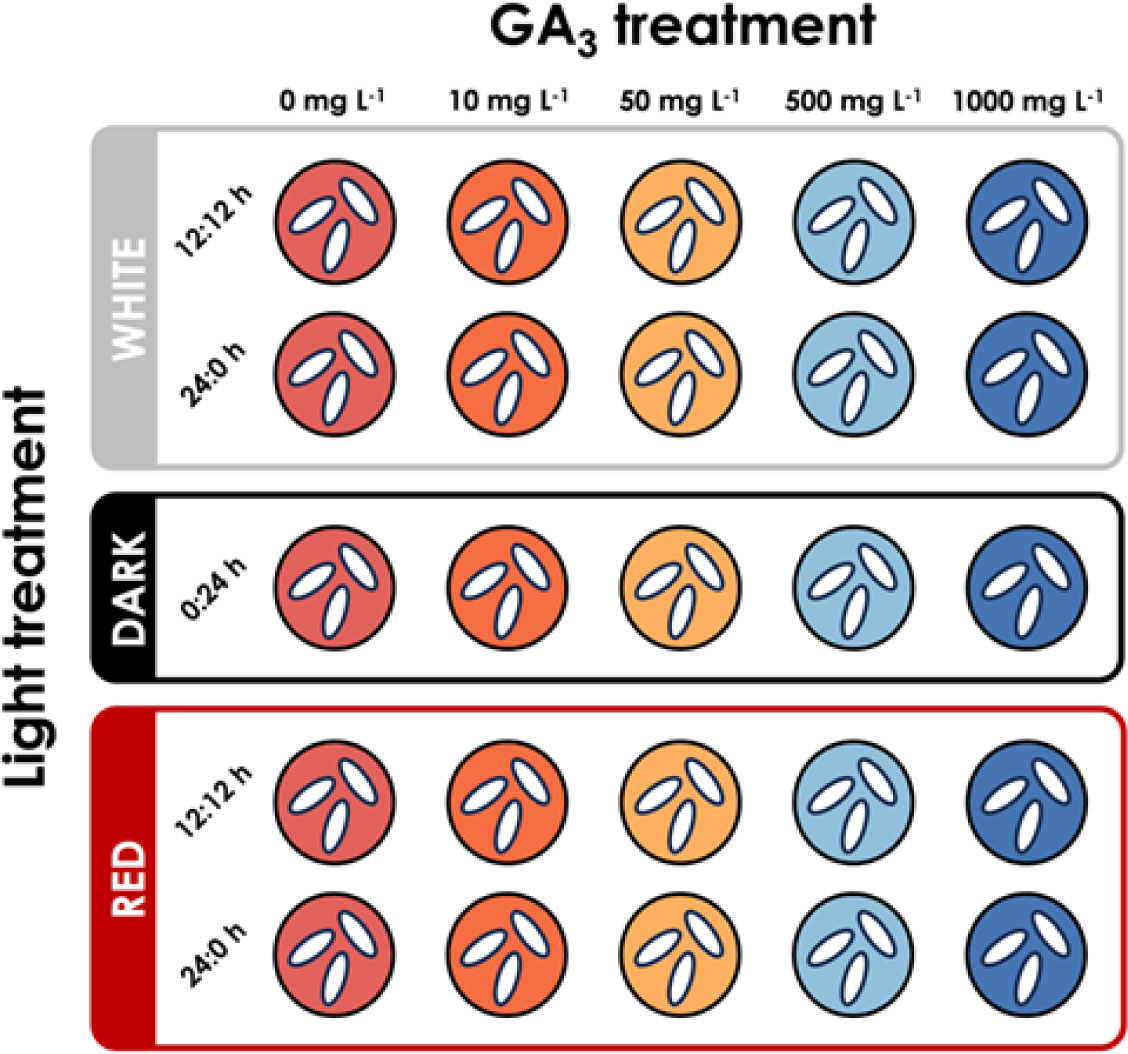
Schematic representation of the experimental set-up. Three seeds of *Zostera marina* were placed in a single petri dish, supplemented with four different concentrations of GA_3_ and a negative control without phytohormone. The petri dishes were exposed to three light spectra and photoperiods. Each petri dish was replicated five times.

Incubation occurred under three light conditions or spectra: white light (400 – 720 nm, 5000 K, 158 μmol m^-2^ s^-1^, BlueView), darkness (no light), and red light (620 – 750 nm, intensity 144 μmol m^-2^ s^-1^ BlueView). We considered white and red light with photoperiods of 12:12h and 24:0h (light:dark regime), as well as a darkness incubation (i.e., 0:24h light:dark regime). Photosynthetically active radiation (PAR) was measured by means of a handheld full-spectrum quantum metre (MQ-200, Apogee).

Each treatment, defined by the combination of light spectrum, photoperiod, and phytohormone, was replicated five times. Within each treatment, replication occurred three times, with three seeds in each experimental unit, resulting in a total of 375 seeds (Fig. **1**).

Experimental units, consisting of petri dishes (35 mm, 5 mL, Avantor), were incubated at 10 ± 0.5°C for the entire duration of the experiment. The reference factor (i.e., “control”) was assumed to be white light, 12:12h photoperiod, and GA_3_ 0 mg L^-1^ (only SSW).

### Germination assessment and seedling development measurement

Germination was assessed three times a week for a total of 35 days. Seed germination was defined as the emergence of the cotyledon from the seed coat (Coolidge, 1983; Liu *et al*., 2023). Culture media (GA_3_ + SSW) were refreshed weekly.

Germinated seeds (hereafter referred to as seedlings) were transferred to a new experimental unit (petri dish, 50 mm, 10 mL, Avantor) and supplemented with SSW (phytohormone-free media).

Seedlings were incubated at 10 ± 0.5 °C under white light at a mean intensity of 158 μmol m^-2^ s^-1^ with a 12:12h photoperiod in a phytohormone-free medium containing SSW. SSW was refreshed weekly to maintain the same nutrient composition. A picture of each seedling was taken twice a week using a microscope (Leica MZ16) coupled with a Canon EOS 600D camera. Pictures were analysed using the ImageJ version 1.52d software for morphometric measurement of the seedlings (Schneider *et al*., 2012). A certified micrometre reticle (Pyser-SGI, serial n. SC1539) was used to calibrate the software for pixel-μm conversion. Morphometric measurements were recorded for each seedling, cotyledon, number of roots, and true leaves, with each of their lengths measured two times a week (Fig. **S1)**. True leaves were defined as the leaves capable of performing photosynthesis, as described by Sugiura *et al*. (2009) and Taylor (2011). The length of the cotyledon was defined as the distance from the basal hypocotyl to the tip of the cotyledonary blade, and the length of true leaves was defined as the distance from the base of the cotyledonary sheath to the tip of the leaf (Fig. **S1**). Maximum growth was calculated by summing the linear measurements of cotyledon and leaf length.

### Statistical analysis

Results are presented as arithmetic mean ± SE, unless stated otherwise. All analyses were performed using R statistical software (v.4.0.2; R Core Team, 2020).

Germination percentage (GP) is the percentage of the cumulative number of germinated seeds at the end of the experiment:

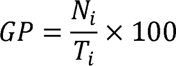

where **T_i_** is the total number of seeds and **N*_i_*** is the number of germinated seeds of a specific treatment.

The effect of light spectrum, photoperiod, and phytohormone concentration on germination was analysed using a penalised logistic regression model implemented through the R package *logistf* (Firth, 1993; Heinze & Puhr, 2010; Puhr *et al*., 2017). Firth’s penalisation approach was selected to account for low event rates and mitigate bias in coefficient estimates. Backward stepwise selection between models was carried out using the Akaike Information Criterion (AIC). ANOVAs, and Tukey’s HSD tests were performed to compare nested models and determine the exclusion of a variable. The outcome variable, germination probability (GB), represents the likelihood of seed germination under varying conditions of light, photoperiod, and phytohormone concentration.

To investigate the effect of light and phytohormone priming on the time to germination, a time-to-event (survival) approach was used via the R package *survival* and *survimer* (Therneau & Grambsch, 2000). Survival analysis considers not only the occurrence of an event, such as germination (binary response), but also the duration it takes for the event (germination) to occur (as a continuous response). Survival model used a Cox proportional hazards method, allowing it to account for both categorical and continuous predictors (McNair *et al*., 2012), with predictors being light, spectrum, and phytohormone and the response variable being time to germination (days). To ensure accurate comparison between the statistical analyses (penalized logistic regression and Cox proportional hazard model), the selection of variables was based on the outcomes of the logistic model’s ANOVA. To visualise the effect of predictors on the response variable, survival curves were computed via the Kaplan-Meier method through the R package *survival*.

Two-way ANOVAs were used to investigate the effect of priming treatments on cotyledon and leaf growth after 60 days, with light and GA_3_ used as categorical explanatory variables. Normality of residuals and homogeneity of variances were verified using Shapiro-Wilk (Shapiro.test, *Stats* package) and Levene’s tests (LeveneTest, *Stats* package), respectively. Leaf length measurements were sqrt-transformed to assure normality of residuals and homogeneity of variances.

## Results

### Influence of priming treatment on germination percentage and germination probability

Overall, GP was 11.67 ± 2.07%, with a daily germination of 0.80 ± 0.40 % seeds. Seeds primed under red light spectrum (combining all GA_3_ treatments) exhibited an overall higher total GP compared to other light treatments (53.57 ± 1.80%) including darkness. Nevertheless, the highest GP was observed in seeds exposed to no light and 10 mg L^-1^ GA_3_ (26.67 ± 2.29%, Fig. **2a**).

**Fig. 2.**
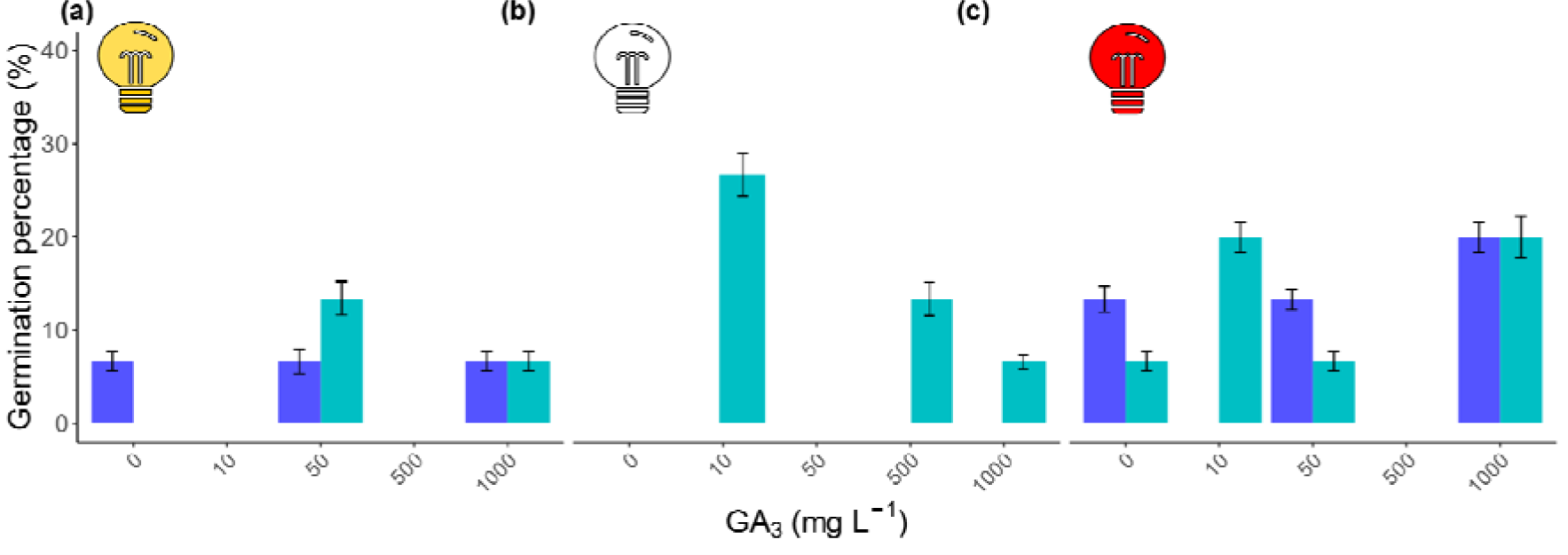
Germination percentage (GP) across the different light, photoperiod and GA_3_ priming treatments. Light bulbs represent the light regime seeds were exposed (yellow bulb: white light; empty bulb: darkness; red bulb: red light). Barplots display the result with standard error bars. Blue barplots represent a 12:12h photoperiod, whereas aquamarine barplots represent a 0:24h (light:dark) photoperiod

Darkness positively affected germination, where a negative dose-effect trend was observed with increasing concentrations of GA_3_ (Fig. **2b**). Moreover, when seeds were incubated in the absence of GA_3_ and in darkness condition, no germination was observed.

Notably, when a specific light spectrum or darkness was combined with a specific concentration of GA_3_, especially at mid-doses of GA_3_ (e.g., no light and 50 mg L^-1^ GA_3_), it often resulted in no germination. This suggests that the presence a specific combination of light condition with doses of GA_3_ can nullify the germination-promoting effect of GA_3_.

After variable selection, the final logistic model included the factors phytohormone, light, and day, but not photoperiod, as well as the interaction of phytohormone and day, and phytohormone and light.

GB was significantly enhanced in seeds primed with either GA_3_ (χ²(4) 42.55, p = 1.28 e^-08^), light and GA_3_ (χ²(8) 114.66, p < 0.001), as well as the extended exposure to GA_3_ overtime (χ²(4) 14.76, p = 5.21 e^-03^).

Penalised logistic regression detected a positive effect of 500 mg L^-1^ GA_3_ (z = 3.42, p = 1.56 e^-06^) and 1000 mg L^-1^ GA_3_ (z = 2.02, p = 2.30 e^-06^) on the germination probability, respectively of 99.86 ± 19.36% and 98.24 ± 19.92%, when the variable time (e.g., day) was held constant.

White light significantly affected GB only when seeds were exposed to 50 mg L^-1^ GA_3_ (z = 3.41, p = 1.59 e^-06^) or 1000 mg L^-1^ GA_3_ (z = 2.02, p = 2.30 e^-02^) (Fig. **S2a**).

A synergistic effect of red light and GA_3_ 10 mg L^-1^ (z = 1.95, p = 1.21 e ^-04^) was observed with a GB 95.4 ± 15.5% compared to control treatments. Further, GB was significantly higher in seeds exposed to darkness at low (z = 3.08, p = 8.18 e^-06^) and mid (z = 2.71, p = 1.68 e^-04^) GA_3_ levels, of respectively 99.8 ± 20.5% for 10 mg L^-1^ GA_3_ and 99.6 ± 20.4% for 500 mg L^-1^ GA_3_ (Fig. **S2b**).

Nevertheless, increasing GA_3_ levels did not consistently influence GB across treatments, with optimal concentrations not exhibiting a dose-dependent response. Notably, seeds exposed to specific concentrations of GA_3_ and light spectrums displayed a drop in GB; for example, when red spectrum was combined with 500 mg L^-1^ GA_3_ (Fig. **S2c**), or darkness with 50 mg L^-1^ GA_3_ (Fig. **S2b**), or white spectrum with 10 and 500 mg L^-1^ GA_3_ (Fig. **S1a**).

### Temporal patterns of seed germination: a time-to-event approach

Overall, for seeds primed with GA_3_ (white spectrum), the likelihood of germination increased 2.72 ± 0.61 times for seeds exposed to 1000 mg LC¹ GA_3_, 1.96 ± 0.64 times for seeds exposed to 50 mg LC¹ GA_3_, and 1.74 ± 0.63 times for seeds exposed to 10 mg LC¹ GA_3_. Seeds exposed to white light and 50 mg LC¹ GA_3_ exhibited the fastest germination response, taking 6.67 ± 1.26 days on average (Fig. **3a**). Notably, only the combination of the red spectrum and 500 mg LC¹ GA_3_ significantly reduced the time to germination (p = 0.033, Fig. **3c**). According to the Cox proportional hazard model, seeds primed under red light germinated 1.83 ± 0.48 times faster as compared to those primed under white light, though this difference was not statistically significant (p = 0.09).

**Fig. 3.**
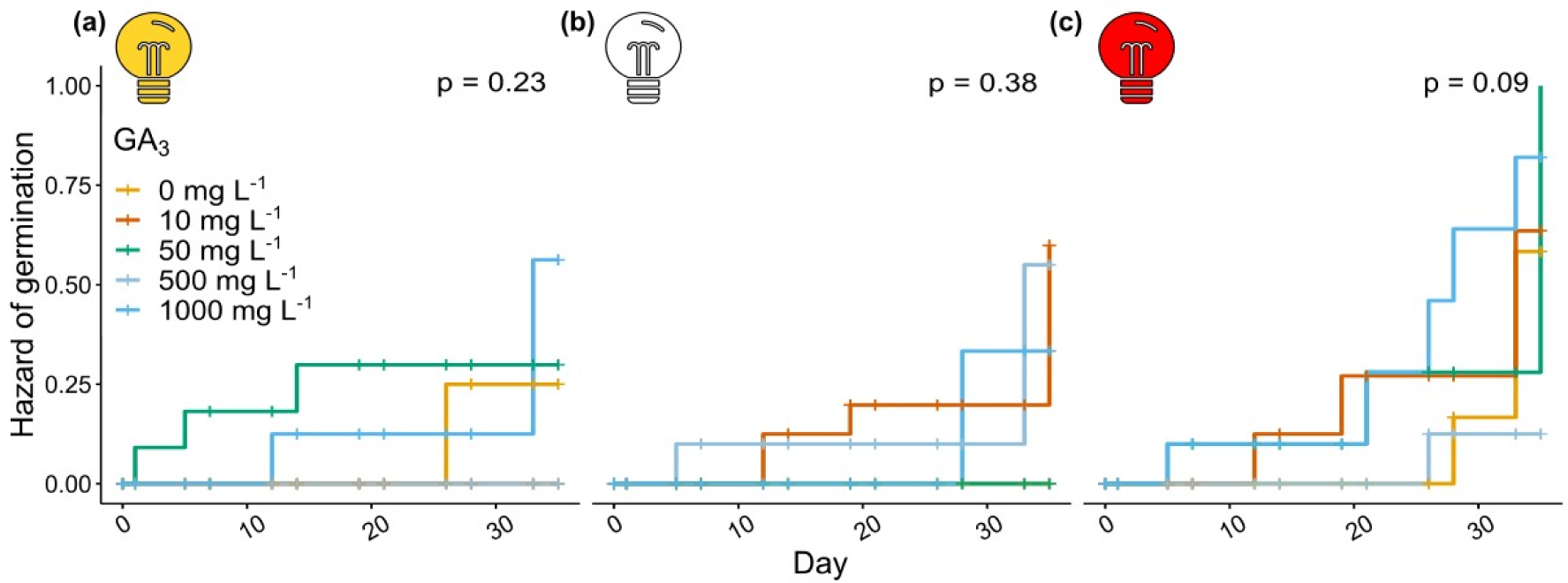
Kaplan-Meier survival curves display the likelihood of experiencing germination across the different light and GA3 priming treatments. Light bulbs represent the light regime seeds were exposed (yellow bulb = white light, empty bulb = darkness and red bulb = red light). P-values indicate whether each light priming treatment have the same germination probability over time.

### Seedling development

The *a posteriori* effect of seed priming on seedling development was monitored to assess potential side-effects on seedling growth. After 60 days, cotyledons reached a mean maximum length of 18.96 ± 5.06 mm and grew at a mean rate of 0.75 ± 0.21 mm day^-1^, while the mean maximum length was 16.78 ± 9.21 mm with a growth rate of 0.84 ± 0.39 mm day^-1^. The first true leaf appeared on average 10.9 ± 2.80 days after germination.

Maximum cotyledon length (19.41 mm) was recorded for seeds incubated in darkness and 10 mg L^-1^ GA_3_ (Fig. **4b**), while maximum leaf length (34.01 mm) was recorded for seeds incubated under red light and 10 mg L^-1^ GA_3_ (Fig. **4f**).

**Fig. 4.**
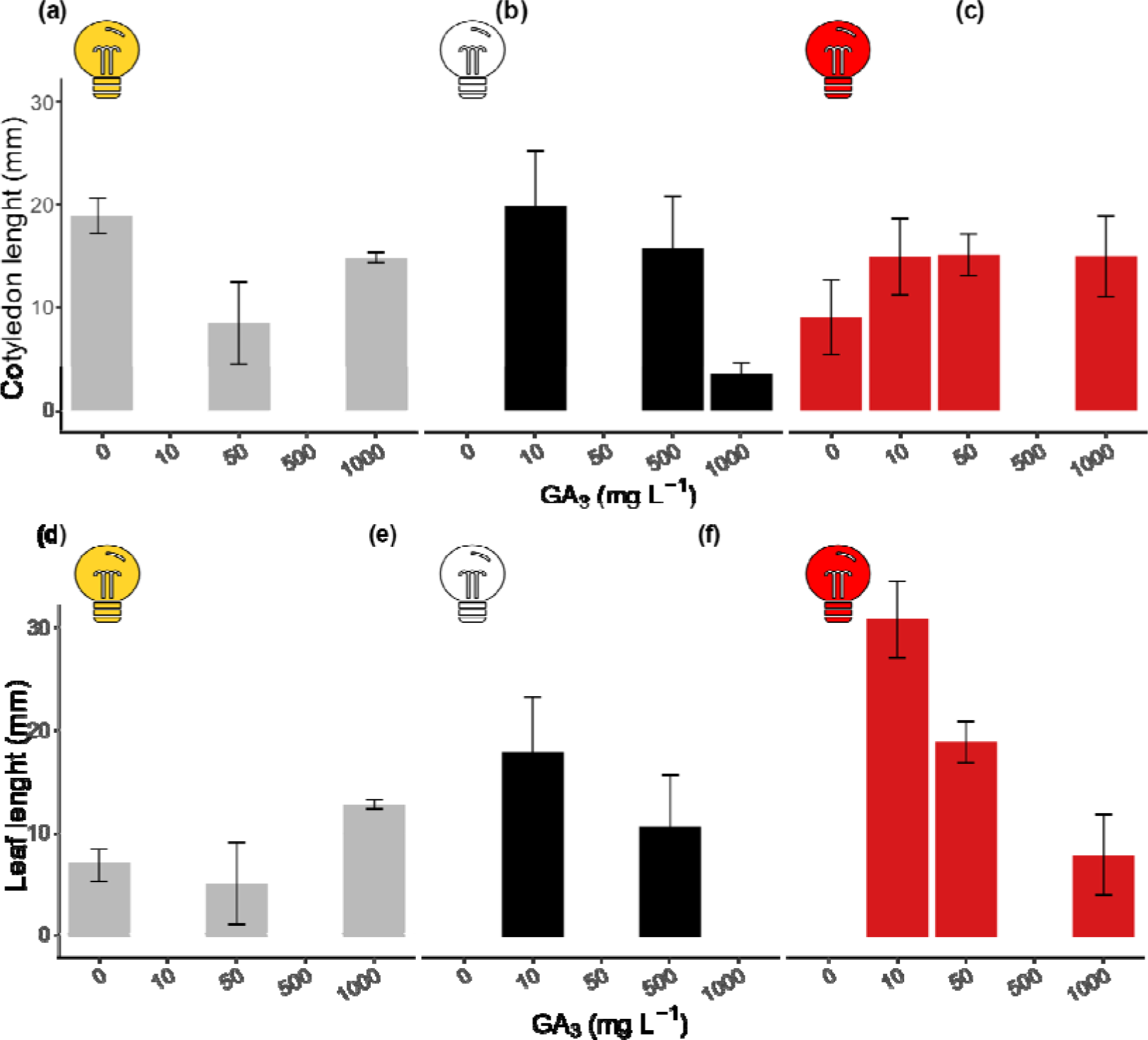
Seedling development assessed through morphometric measurements of a) Cotyledon and b) Leaf length (mm) of the germinated primed seeds. Seedlings were incubated in SSW under white light. Light and GA_3_ priming are indicated to show the applied treatments. Light bulbs represent the light regime seeds were exposed (yellow bulb = white light, empty bulb = darkness and red bulb = red light). Barplots display the result with standard error bars.

We found no effect of priming treatment on cotyledon length (p = 0.17) and leaf length (p = 0.08).

## Discussion

Our findings provide new experimental evidence that light conditions and GA_3_ priming increase germination rates of seeds from the seagrass *Zostera marina*, likely by influencing the physiological processes conducive to germination without affecting the subsequent stages of seedling development.

### Effects of light on germination

Seed germination under light priming suggests an active involvement of the plant photoreceptors in the germination process (Mérai *et al*., 2019). In *Z. marina*, a total of six photoreceptors have been identified, each potentially contributing to different signalling pathways associated with the regulation of seed germination and enzymatic processes weakening the seed walls (Olsen *et al*., 2016).

In this study, we observed that seeds exposed to white light, which encompasses both photosynthetically active red and blue wavelengths, displayed inhibited germination compared to seeds solely exposed to red light. This reduced germination under white light may be attributed to the activation of cryptochromes by the blue component of the light spectrum (Jacobsen *et al*., 2013; Barrero *et al*., 2014; Hofmann, 2014), which enhances embryo sensitivity by upregulating ABA-associated genes (Hoang *et al*., 2014). This phenomenon can be explained by considering the natural habitat of intertidal *Z. marina* meadows, where seeds covered by a sediment layer can still be exposed to daylight, experiencing a shifted R:FR ratio and reduced irradiance. Similarly, seeds covered by a layer of water during low tide encounter light that shifts towards more blue and green wavelengths.

Photoinhibition of germination following white light exposure has been documented for several terrestrial angiosperms and is interpreted as an adaptive trait to avert germination in suboptimal conditions found in extreme and dynamic environment ecosystems (e.g., coastal dunes, deserts) (Mérai *et al*., 2023). We therefore speculate that this form of photoinhibition may also occur in highly dynamic marine environments, such as intertidal ecosystems and shallow waters, where temperature, salinity, and irradiance can vary greatly over a short period of time. Moreover, although our results do not indicate a significant effect of the photoperiod on germination under white light, it is believed that inhibited germination following exposure to white light is a result of a photoperiod measuring mechanism for optimising embryo development (Mérai *et al*., 2023).

In *Z. marina,* a total of six photoreceptors have been identified, each potentially contributing to diverse signalling pathways associated with the regulation of seed germination and enzymatic processes that weaken the seed walls (Mawphlang & Kharshiing, 2017; Nautiyal *et al*., 2023). Moreover, *Z. marina* appears to have lost a number of photoreceptors (cry2, cry5, cryDASH and phyC) during the evolutionary adaptation towards the light-attenuated habitat of the marine environment; however, it maintained cry1 (cryptochrome), phot1, and phot2 (phototropins), which are likely involved in transducing blue light signals, as well as phyA and phyB (phytochromes), which mediate red light transduction.

Red-light mediated germination is reported for many terrestrial (Mathews, 2006) and aquatic floating species (Ziegler *et al*., 2023), where the phytochrome inactive form Pr converts to the active Pfr upon absorption of red light and initiates the signalling cascades that ultimately lead to germination (Furuya & Schäfer, 1996). Wang *et al*. (2017) provided the first evidence on the effect of light quality on the germination of *Z. marina* seeds in Lidao Bay (China), observing that red light has no effect on germination. The present research built upon the previous knowledge with a specialised set-up considering a reduced photosynthetically active radiation (light intensity), salinity, and temperature, conditions more commonly found in the shallow spring waters in the North Atlantic region. Strikingly, in our study, we observed improved germination when seeds were exposed to red light. We attribute these different outcomes to potential population-specific adaptations to red light or as an adaptation to sustain germination during seasonal increases in suspended organic matter, which may shift irradiance in the water column towards yellow-red wavelengths (Strydom *et al*., 2017). Evidence of the positive effects of red light on germination has been reported for several other seagrass species, such as *H. ovalis* (Waite *et al.,* 2021), *Thalassia hemprichii* (Soong *et al.,* 2013b) and *Z. japonica* (Kaldy *et al.,* 2014).

### Effects of GA_3_ priming on germination

GA_3_ seed priming is a process used to initiate a pre-germinative metabolic state, enhancing the germination and seed vigour of many terrestrial species of commercial (Kaur *et al*., 2006; Lee *et al*., 2018; Azab, 2018; Budzeń *et al*., 2018), as well as conservation interest (Yang *et al*., 2005; Barden *et al*., 2017). Further, GA_3_ is widely employed in horticulture to stimulate germination and improve, for instance, the size and quality of fruits (Rademacher, 2015).

Similar to terrestrial plants, in our study, *Z. marina* seeds exposed to GA_3_ displayed a greater germination percentage as compared to non-primed seeds. However, no studies have reported the use of GA_3_ as an exogenous hormone treatment to stimulate the germination in *Z. marina* seeds; therefore, to identify optimal and effective concentrations of GA_3_ a broader range of concentrations was adopted. In *Posidonia australis*, Glasby *et al*. (2015) found no effect of 0.1 mg L^-1^ and 20 mg L^-1^ GA_3_ on germination. Similarly, Loques *et al*. (1990) observed that 1 mg L^-1^ GA_7_ had no effect on germination in *Zostera noltii*. However, the highly saline media employed in their experiment may have acted as a suppressor of germination during the limited experimental time (Hootsmans *et al*., 1987). Indeed, our earliest germination was recorded in a seed primed with 10 mg L^-1^ GA_3_ and white light after 24 hours, suggesting the efficacy of the treatment to be strongly time-dependent. Conacher *et al*. (1994) found that germination in *Zostera capricorni* was promoted by a treatment of 50 mg L^-1^ GA_3_ but not by 500 mg L^-1^ GA_3_; similar results were also observed in our study, except for seeds under darkness conditions. This discrepancy may be explained by a compensatory mechanism adopted by the seed embryo in the physiological regulation after the exposure to light and/or hormone. For instance, a study conducted by Toyomasu *et al*. (1998) demonstrated, on a terrestrial species, that red light germination requirement can be mimicked by exposing the seeds to a concentration of 34.64 mg L^-1^ GA_3_ (10^−4^ M), with the same germination outcomes between treated and control seeds. Hence, light spectrum and hormones can synergistically interact in the regulation of germination (Shinomura, 1997), but they can also antagonistically regulate germination, neutralising each other’s effect. Overall, the responses of *Z. marina*, *Z. capricorni*, and *Z. noltii* seeds to GA_3_ treatments may also be species-specific, since the genetic and molecular mechanisms controlling germination differ considerably, even between populations of the same seagrass species (Xu *et al*., 2021; Zhang *et al*., 2022). Additionally, changes at the level of seagrass genomes over the course of their adaptation to the marine environment include changes in hormone signalling and biosynthesis (Chen *et al*., 2021). The GAs signalling pathway appears to have been modified in *Z. muelleri* (Lee *et al*., 2016) and in *Z. marina* with the loss of genes associated with ethylene biosynthesis (Golicz *et al*., 2015). Nevertheless, the roles of individual genes in the GA signalling pathway in seagrasses are, yet, poorly understood (Chen & Qiu, 2022), and further understanding of the genes involved in the mechanisms regulating germination could shed light on associated hormonal pathways and their response to hormone treatments.

While our study provides valuable insights into the combined effects of light and phytohormones on seed germination and dormancy in *Z. marina*, it is important to critically assess our findings, particularly some unexpected results where specific combinations of light spectra and phytohormones resulted in enhanced dormancy. GA_3_ appeared to inhibit germination at specific concentrations, with 500 mg L^-1^ being the most recurrent one, especially under white and red light incubation, but not in darkness. For some terrestrial species, evidence suggests that light mediates the modulation of GA, altering seed sensitivity during germination to exogenous GAs. Studies on *Arabidopsis* revealed that white light-primed seeds did germinate at lower exogenous concentrations of GA (Yang *et al*., 1995), while exposure of seeds to different red and far-red (R:FR) ratios, under red light, enhanced GA signalling (Poppe & Schäfer, 1997; Halliday & Fankhauser, 2003). Similarly, germination inhibition was observed at 10 and 50 mg L^-1^ for white light and darkness, respectively.

Moreover, these germination no-responses challenge traditional dose-response assumptions and suggest a non-monotonic relationship between GA_3_ concentrations and seed germination. Several explanations were considered to understand this phenomenon, including a non-monotonic dose-response (Cvrčková *et al*., 2015), complex hormonal interactions and modulation (Vanstraelen & Benková, 2012), physiological thresholds (Bradford & Trewavas, 1994; Fennimore & Foley, 1998), seed sensitivity to GA_3_ (Jalal *et al*., 2021), and potential antagonistic effects of specific GA_3_ concentrations when combined with light (Seo *et al*., 2009), and experimental limitation (e.g., total number of seeds, limited replication). The lack of germination at certain GA_3_ concentrations warrants further investigation. We, therefore, suggest exploring intermediate GA_3_ concentrations and incorporating gene expression analysis and hormonal profiling. These results underscore the complexity of hormonal regulation in plants, and more specifically in this marine seagrass species, *Z. marina*.

### Impact of seed priming on early-stage seedling development

Although the present research did not expose *Z. marina* seedlings to any of the priming agents tested, we cannot disregard the potential backdrop effects of the seed priming on their later development. Indeed, GA_3_ and light priming are often adopted in terrestrial plants to promote and support seedling development (Rademacher, 2015; Waqas *et al*., 2019), with, for instance, GA_3_ enhancing primary and secondary growth in adult plants of sugar beet and korarima (Eyob, 2009; Azab, 2018). A similar approach has been studied for some marine plant species, with, for instance, GA_3_ in concentrations ranging from 0.01 to 35 mg L^-1^ enhancing leaf and rhizome growth in *Cymodocea nodosa* (Zarranz *et al*., 2010). However, such effect of GA_3_ was not reported for *Posidonia australis* (Glasby *et al*., 2015). For *Zostera noltii*, Loques *et al*. (1990) found that 1 mg L^-1^ GA_7_ negatively affected seedling growth, while in *H. ovalis* light quality was responsible for reduced above- and below- ground biomass (Strydom *et al*., 2017).

Overall, our study found that no priming treatment negatively affected the growth, morphology, and survival of cotyledon and leaves in seedlings. Subsequent seedling growth rates were observed to be comparable to previously reported values where no treatment was adopted (Taylor, 1957; van Lent & Verschnure, 1995; Niu *et al*., 2012). While our results suggest that seed priming does not produce adverse effects on the development of *Z. marina* seedlings, it is plausible that our findings provide only a partial understanding of the potential effects of seed priming on seedling development, especially considering that some GA_3_ concentrations did not deliver germinative outcomes.

In future studies, we recommend combining morphometric measurements with assessments of the photosynthetic responses of the seedlings and leaf pigment analysis. This integrated approach would offer a more comprehensive understanding of the effects of seed priming on *Z. marina* development.

### Implications for restoration

Seagrass conservation and restoration are critical priorities emphasized by the United Nations (UNEP, 2020) and the European Union (COM, 2024). In response to climate change impacts on ecosystems, seed-based restoration of *Z. marina* habitats is being implemented to restore this widely distributed seagrass species. The seagrass community is intensifying efforts to advance both scientific and logistical methods and standards for scaling up seagrass restoration. Enhancing the success and maximising the utilization of seed stocks will strengthen restoration initiatives and support ecosystem resilience (Nordlund *et al*., 2024).

To ensure the affordability and scalability of restoration projects, it is important for practitioners to explore cost-effective methods that enhance their efforts. Over the past decades, researchers have described and improved seed-based restoration techniques (Unsworth *et al*., 2019a; Tan *et al*., 2020; Gräfnings *et al*., 2023), methods to support germination (Waite *et al*., 2021; Xu *et al*., 2021), and seedling establishment (Verduin *et al*., 2013). Previous techniques, such as seed scarification and stratification, have yielded mixed and unpredictable outcomes. Enhancing seed germination rates and optimising nursery practices are critical steps in overcoming current limitations of seed availability and meeting the escalating demand for restoration efforts.

Our study shows that seed priming, involving treatments with light and phytohormones, presents a promising approach to increase seed germination potential before sowing. This research represents a first attempt to showcase the potential feasibility of light and hormone seed priming to promote germination and release dormancy in contexts where on-demand germination is required, such as nurseries or restoration projects. Further studies should explore the optimal concentrations and incubation durations required to induce the pre-germinative stage (early stages of germination) without triggering the actual sprouting prior sowing. This approach has the potential to enhance germination rates, reduce time to germination, and achieve more uniform germination once the seeds are sown.

Moreover, as this study is conducted in a laboratory setting, we acknowledge the potential limitations of our study, which may overlook variables or interactions that are at play in real-life conditions, thus distancing our outcomes from full practical application. We therefore support and promote the transfer of this lab-based knowledge to field settings to achieve a more complete understanding of the efficacy of priming methods in natural environments, striving towards successful seagrass restoration projects.

## Conclusion

The modulation of seed dormancy and germination through photo-hormonal regulation plays a pivotal role within the life cycles of both terrestrial and aquatic plants. This study demonstrated the positive effect of light and GA_3_ priming on seed germination of *Z. marina* by investigating the intricate germination responses associated with seed physiology following exposure to priming agents. The results indicate that light and phytohormone priming can act as a catalytic process, inducing germination and releasing dormancy following seed manipulation or overwintering. Red light spectrum and darkness when combined with GA_3_ positively induce germination; however, these responses are more likely to follow a non-monotonic dose response. Furthermore, seed priming did not produce *a posteriori* negative effects on seedling growth, showcasing the potential feasibility of the application of seagrass seed priming in other practices where on-demand germination is required. (i.e., seagrass nursery, seed-based restoration project).

## Acknowledgements

This research was supported by Flanders innovation & entrepreneurship (VLAIO) and the Research Foundation Flanders (FWO). The research leading to results presented in this publication was carried out with infrastructure funded by EMBRC Belgium - FWO international research infrastructure I001621N.

We gratefully thank the Alfred Wegener Institute and especially, Landesamt für Umwelt des Landes Schleswig-Holstein (LfU), and Landesbetrieb für Küstenschutz, Nationalpark und Meeresschutz Schleswig-Holstein, Nationalparkverwaltung for the local guidance, supporting the permit request.

## Competing interest

None.

## Author contributions

RP planned and designed the research. TD requested the sampling permit to the competent authority. RP and LW performed experiments and analysed the data. RP and TVdS prepared the figures. RP led the writing of the manuscript, with extensive input from TVdS, NK and AV.

## Data availability

The data that support the findings of this study are available from the corresponding author upon request.

## Notes

### Competing Interest Statement

The authors have declared no competing interest.

## References

Azab E. 2018. Seed Pre-soaking on Gibberellic Acid (GA3) Enhance Growth, Histological and Physiological Traits of Sugar Beet (Beta vulgaris L) under Water Stress. Egyptian Journal of Agronomy 40: 119–132.

Barden C, Boyer C, Morales B, Fisher L. 2017. Promoting Red Elm (*Ulmus rubra* Muhl.) Germination with Gibberellic Acid. Journal of Forestry 115: 393–396.

Barrero JM, Downie AB, Xu Q, Gubler F. 2014. A Role for Barley CRYPTOCHROME1 in Light Regulation of Grain Dormancy and Germination. The Plant Cell 26: 1094–1104.

Bradford KJ, Trewavas AJ. 1994. Sensitivity Thresholds and Variable Time Scales in Plant Hormone Action. Plant Physiology 105: 1029–1036.

Budzeń M, Kornacki A, Kornarzyński K, Sujak A. 2018. Effect of laser light on germination of *Lavatera thuringiaca* L. seeds studied by binomial distribution. Seed Science and Technology 46: 447–464.

Chen S, Qiu G. 2022. Overexpression of the intertidal seagrass J protein ZjDjB1 enhances tolerance to chilling injury. Plant Biotechnology Reports 16: 419–435.

Chen J, Zang Y, Shang S, Liang S, Zhu M, Wang Y, Tang X. 2021. Comparative Chloroplast Genomes of Zosteraceae Species Provide Adaptive Evolution Insights Into Seagrass. Frontiers in Plant Science 12: 741152.

Chua M, Erickson TE, Merritt DJ, Chilton AM, Ooi MKJ, Muñoz-Rojas M. 2020. Bio-priming seeds with cyanobacteria: effects on native plant growth and soil properties. Restoration Ecology 28: S168–S176.

Conacher CA, Poiner IR, Butler J, Pun S, Tree DJ. 1994. Germination, storage and viability testing of seeds of *Zostera capricorni* Aschers. from a tropical bay in Australia. Aquatic Botany 49: 47–58.

Cvrčková F, Luštinec J, Žárský V. 2015. Complex, non-monotonic dose-response curves with multiple maxima: Do we (ever) sample densely enough? Plant Signaling & Behavior 10: e1062198.

Dooley FD, Wyllie-Echeverria S, Van Volkenburgh E. 2013. Long-term seed storage and viability of *Zostera marina*. Aquatic Botany 111: 130–134.

Duarte CM. 2002. The future of seagrass meadows. Environmental Conservation 29: 192–206.

Dutta P. 2018. Seed Priming: New Vistas and Contemporary Perspectives. In: Rakshit A, Singh HB, eds. Advances in Seed Priming. Singapore: Springer, 3–22.

Environment UN. 2020. Out of the Blue: The Value of Seagrasses to the Environment and to People. UNEP(2020). Nairob*i*.

European Commission. 2024. Report from the Commission to the European Parliament, the Council and the European Economic and Social Committee: The state of nature in the European Union, Report on the status and trends in 2013 - 2018 of species and habitat types protected by the Birds and Habitats Directives. COM(2024) 635 final (Publications Office of the European Union, 2024)

Eyob S. 2009. Promotion of seed germination, subsequent seedling growth and in vitro propagation of korarima (*Aframomum corrorima* (Braun) P.C.M. Jansen). Journal of Medicinal Plants Research.

Fennimore SA, Foley ME. 1998. Genetic and physiological evidence for the role of gibberellic acid in the germination of dormant *Avena fatua* seeds. Journal of Experimental Botany 49: 89–94.

Finch-Savage WE, Leubner-Metzger G. 2006. Seed dormancy and the control of germination. New Phytologist 171: 501–523.

Firth D. 1993. Bias reduction of maximum likelihood estimates. Biometrika 80: 27–38.

Glasby TM, Taylor SL, Housefield GP. 2015a. Factors influencing the growth of seagrass seedlings: A case study of *Posidonia australis*. Aquatic Botany 120: 251–259.

Glasby TM, Taylor SL, Housefield GP. 2015b. Factors influencing the growth of seagrass seedlings: A case study of *Posidonia australis*. Aquatic Botany 120: 251–259.

Golicz AA, Schliep M, Lee HT, Larkum AWD, Dolferus R, Batley J, Chan C-KK, Sablok G, Ralph PJ, Edwards D. 2015. Genome-wide survey of the seagrass *Zostera muelleri* suggests modification of the ethylene signalling network. Journal of Experimental Botany 66: 1489–1498.

Govers LL, Van Der Zee EM, Meffert JP, Van Rijswick PCJ, Man In T Veld WA, Heusinkveld JHT, Van Der Heide T. 2017. Copper treatment during storage reduces *Phytophthora* and *Halophytophythora* infection of *Zostera marina* seeds used for restoration. Scientific Reports 7: 1–8.

Gräfnings MLE, Heusinkveld JHT, Hoeijmakers DJJ, Smeele Q, Wiersema H, Zwarts M, van der Heide T, Govers LL. 2023. Optimizing seed injection as a seagrass restoration method. Restoration Ecology 31: e13851.

Halliday KJ, Fankhauser C. 2003. Phytochrome-hormonal signalling networks. New Phytologist 157: 449–463.

Heinze G, Puhr R. 2010. Bias-reduced and separation-proof conditional logistic regression with small or sparse data sets. Statistics in Medicine 29: 770–777.

Hoang HH, Sechet J, Bailly C, Leymarie J, Corbineau F. 2014. Inhibition of germination of dormant barley (*Hordeum vulgare* L.) grains by blue light as related to oxygen and hormonal regulation. Plant, Cell & Environment 37: 1393–1403.

Hofmann N. 2014. Cryptochromes and Seed Dormancy: The Molecular Mechanism of Blue Light Inhibition of Grain Germination. The Plant Cell 26: 846.

Hootsmans MJM, Vermaat JE, Van Vierssen W. 1987. Seed-bank development, germination and early seedling survival of two seagrass species from The Netherlands: *Zostera marina* L. and *Zostera noltii* hornem. Aquatic Botany 28: 275–285.

Infantes E, Eriander L, Moksnes P. 2016. Eelgrass (*Zostera marina*) restoration on the west coast of Sweden using seeds. Marine Ecology Progress Series 546: 31–45.

Jacob C, Buffard A, Pioch S, Thorin S. 2018. Marine ecosystem restoration and biodiversity offset. Ecological Engineering 120: 585–594.

Jacobsen JV, Barrero JM, Hughes T, Julkowska M, Taylor JM, Xu Q, Gubler F. 2013. Roles for blue light, jasmonate and nitric oxide in the regulation of dormancy and germination in wheat grain (*Triticum aestivum* L.). Planta 238: 121–138.

Jalal A, Oliveira Junior JC de, Ribeiro JS, Fernandes GC, Mariano GG, Trindade VDR, Reis AR dos. 2021. Hormesis in plants: Physiological and biochemical responses. Ecotoxicology and Environmental Safety 207: 111225.

Jarvis JC, Moore KA. 2010. The role of seedlings and seed bank viability in the recovery of Chesapeake Bay, USA, *Zostera marina* populations following a large-scale decline. Hydrobiologia 649: 55–68.

Kaldy JE. 2014. Effect of temperature and nutrient manipulations on eelgrass *Zostera marina* L. from the Pacific Northwest, USA. Journal of Experimental Marine Biology and Ecology 453: 108–115.

Kaur R, Sharma N, Kumar K, Sharma DR, Sharma SD. 2006. *In vitro* germination of walnut (*Juglans regia* L.) embryos. Scientia Horticulturae 109: 385–388.

Kettenring KM, Tarsa EE. 2020. Need to Seed? Ecological, Genetic, and Evolutionary Keys to Seed-Based Wetland Restoration. Frontiers in Environmental Science 8.

Koornneef M, Bentsink L, Hilhorst H. 2002. Seed dormancy and germination. Current Opinion in Plant Biology 5: 33–36.

Lee H, Golicz AA, Bayer PE, Jiao Y, Tang H, Paterson AH, Sablok G, Krishnaraj RR, Chan C-KK, Batley J, et al. 2016. The Genome of a Southern Hemisphere Seagrass Species (*Zostera muelleri*). Plant Physiology 172: 272–283.

Lee JW, Jo IH, Kim JU, Hong CE, Kim YC, Kim DH, Park YD. 2018. Improvement of seed dehiscence and germination in ginseng by stratification, gibberellin, and/or kinetin treatments. Horticulture Environment and Biotechnology 59: 293–301.

van Lent F, Verschnure JM. 1995. Comparative study on populations of *Zostera marina* L. (eelgrass): experimental germination and growth. Journal of Experimental Marine Biology and Ecology 185: 77–91.

Liu S, Jiang Z, Wu Y, Zhang X, Huang X. 2023. Combined effects of temperature and burial on seed germination and seedling growth rates of the tropical seagrass *Enhalus acoroides*. Journal of Experimental Marine Biology and Ecology 562: 151881.

Loques F, Caye G, Meinesz A. 1990. Germination in the marine phanerogam *Zostera noltii* Hornemann at Golfe Juan, French Mediterranean. Aquatic Botany 38: 249–260.

Marion SR, Orth RJ. 2010. Innovative Techniques for Large-scale Seagrass Restoration Using Zostera marina (eelgrass) Seeds. Acta Horticulturae 528: 514–526.

Mathews S. 2006. Phytochrome-mediated development in land plants: red light sensing evolves to meet the challenges of changing light environments. Molecular Ecology 15: 3483–3503.

Mawphlang OIL, Kharshiing EV. 2017. Photoreceptor Mediated Plant Growth Responses: Implications for Photoreceptor Engineering toward Improved Performance in Crops. Frontiers in Plant Science 8: 1181.

McDevitt-Irwin J, Iacarella J, Baum J. 2016. Reassessing the nursery role of seagrass habitats from temperate to tropical regions: a meta-analysis. Marine Ecology Progress Series 557: 133–143.

McNair JN, Sunkara A, Frobish D. 2012. How to analyse seed germination data using statistical time-to-event analysis: non-parametric and semi-parametric methods. Seed Science Research 22: 77–95.

Mérai Z, Xu F, Musilek A, Ackerl F, Khalil S, Soto-Jiménez LM, Lalatović K, Klose C, Tarkowská D, Turečková V, et al. 2023. Phytochromes mediate germination inhibition under red, far-red, and white light in *Aethionema arabicum*. Plant Physiology 192: 1584–1602.

Moore KA, Orth RJ, Nowak JF. 1993. Environmental regulation of seed germination in *Zostera marina* L. (eelgrass) in Chesapeake Bay: effects of light, oxygen and sediment burial. Aquatic Botany 45: 79–91.

Nautiyal PC, Sivasubramaniam K, Dadlani M. 2023. Seed Dormancy and Regulation of Germination. In: Dadlani M, Yadava DK, eds. Seed Science and Technology: Biology, Production, Quality. Singapore: Springer Nature, 39–66.

Niu S, Zhang P, Liu J, Guo D, Zhang X. 2012. The effect of temperature on the survival, growth, photosynthesis, and respiration of young seedlings of eelgrass *Zostera marina* L. Aquaculture 350–353: 98–108.

Nordlund LM, Unsworth RKF, Wallner-Hahn S, Ratnarajah L, Beca-Carretero P, Boikova E, Bull JC, Chefaoui RM, de los Santos CB, Gagnon K, et al. 2024. One hundred priority questions for advancing seagrass conservation in Europe. PLANTS, PEOPLE, PLANET 6: 587–603.

Olsen JL, Rouzé P, Verhelst B, Lin Y-C, Bayer T, Collen J, Dattolo E, De Paoli E, Dittami S, Maumus F, et al. 2016. The genome of the seagrass *Zostera marina* reveals angiosperm adaptation to the sea. Nature 530: 331–335.

Orth RJ, Fishman JR, Harwell MC, Marion SR. 2003. Seed-density effects on germination and initial seedling establishment in eelgrass *Zostera marina* in the Chesapeake Bay region. Marine Ecology Progress Series 250: 71–79.

Orth RJ, Harwell MC, Bailey EM, Bartholomew A, Jawad JT, Lombana AV, Moore KA, Rhode JM, Woods HE. 2000. A review of issues in seagrass seed dormancy and germination: implications for conservation and restoration. Marine Ecology Progress Series 200: 277–288.

Orth RJ, Marion SR, Moore KA. 2007. A summary of eelgrass (*Zostera marina*) reproductive biology with an emphasis on seed biology and ecology from the Chesapeake Bay region. US Army Engineer Research and Development Center.

Poppe C, Schäfer E. 1997. Seed Germination of *Arabidopsis thaliana* phyA/phyB Double Mutants Is under Phytochrome Control. Plant Physiology 114: 1487–1492.

Probert RJ, Daws MI, Hay FR. 2009. Ecological correlates of ex situ seed longevity: a comparative study on 195 species. Annals of Botany 104: 57–69.

Puhr R, Heinze G, Nold M, Lusa L, Geroldinger A. 2017. Firth’s logistic regression with rare events: accurate effect estimates and predictions? Statistics in Medicine 36: 2302–2317.

Rademacher W. 2015. Plant Growth Regulators: Backgrounds and Uses in Plant Production. Journal of Plant Growth Regulation 34: 845–872.

Reynolds LK, Waycott M, McGlathery KJ. 2013. Restoration recovers population structure and landscape genetic connectivity in a dispersal-limited ecosystem. Journal of Ecology 101: 1288–1297.

Riddin T, Adams JB. 2009. The seed banks of two temporarily open/closed estuaries in South Africa. Aquatic Botany 90: 328–332.

Schneider CA, Rasband WS, Eliceiri KW. 2012. NIH Image to ImageJ: 25 years of image analysis. Nature Methods 9: 671–675.

Seo M, Nambara E, Choi G, Yamaguchi S. 2009. Interaction of light and hormone signals in germinating seeds. Plant Molecular Biology 69: 463–472.

Serrano O, Gómez-López DI, Sánchez-Valencia L, Acosta-Chaparro A, Navas-Camacho R, González-Corredor J, Salinas C, Masque P, Bernal CA, Marbà N. 2021. Seagrass blue carbon stocks and sequestration rates in the Colombian Caribbean. Scientific Reports 11: 11067.

Shinomura T. 1997. Phytochrome regulation of seed germination. Journal of Plant Research 110: 151–161.

Short F, Carruthers T, Dennison W, Waycott M. 2007a. Global seagrass distribution and diversity: A bioregional model. Journal of Experimental Marine Biology and Ecology 350: 3–20.

Short F, Carruthers T, Dennison W, Waycott M. 2007b. Global seagrass distribution and diversity: A bioregional model. Journal of Experimental Marine Biology and Ecology 350: 3–20.

Sinclair EA, Verduin J, Krauss SL, Hardinge J, Anthony J, Kendrick GA. 2013. A genetic assessment of a successful seagrass meadow (*Posidonia australis*) restoration trial. Ecological Management & Restoration 14: 68–71.

Soong K, Chiu S-T, Chen C-NN. 2013. Novel Seed Adaptations of a Monocotyledon Seagrass in the Wavy Sea (M Bennett, Ed.). PLoS ONE 8: e74143.

Statton J, Sellers R, Dixon KW, Kilminster K, Merritt DJ, Kendrick GA. 2017. Seed dormancy and germination of *Halophila ovalis* mediated by simulated seasonal temperature changes. Estuarine, Coastal and Shelf Science 198: 156–162.

Strydom S, McMahon K, Kendrick G, Statton J, Lavery P. 2017. Seagrass *Halophila ovalis* is affected by light quality across different life history stages. Marine Ecology Progress Series 572: 103–116.

Sugiura H, Hiroe Y, Suzuki T, Maegawa M. 2009. The carbohydrate catabolism of *Zostera marina* influenced by lower salinity during the pre-germination stage. Fisheries Science 75: 1205–1217.

Tan YM, Dalby O, Kendrick GA, Statton J, Sinclair EA, Fraser MW, Macreadie PI, Gillies CL, Coleman RA, Waycott M, et al. 2020. Seagrass Restoration Is Possible: Insights and Lessons From Australia and New Zealand. Frontiers in Marine Science 7: 617.

Taylor ARA. 1957. STUDIES OF THE DEVELOPMENT OF *ZOSTERA MARINA* L.: II. GERMINATION AND SEEDLING DEVELOPMENT. Canadian Journal of Botany 35: 681–695.

Taylor A. 2011. Studies of the development of *Zostera marina* L. I. The embryo and seed. Canadian Journal of Botany 35: 477–499.

Toyomasu T, Kawaide H, Mitsuhashi W, Inoue Y, Kamiya Y. 1998. Phytochrome Regulates Gibberellin Biosynthesis during Germination of Photoblastic Lettuce Seeds. Plant Physiology 118: 1517–1523.

Turner SR, Steadman KJ, Vlahos S, Koch JM, Dixon KW. 2013. Seed Treatment Optimizes Benefits of Seed Bank Storage for Restoration-Ready Seeds: The Feasibility of Prestorage Dormancy Alleviation for Mine-Site Revegetation. Restoration Ecology 21: 186–192.

Unsworth RKF, Bertelli CM, Coals L, Cullen-Unsworth LC, den Haan S, Jones BLH, Rees SR, Thomsen E, Wookey A, Walter B. 2023. Bottlenecks to seed-based seagrass restoration reveal opportunities for improvement. Global Ecology and Conservation 48: e02736.

Unsworth RKF, Bertelli CM, Cullen-Unsworth LC, Esteban N, Jones BL, Lilley R, Lowe C, Nuuttila HK, Rees SC. 2019a. Sowing the Seeds of Seagrass Recovery Using Hessian Bags. Frontiers in Ecology and Evolution 7.

Unsworth RKF, McKenzie LJ, Collier CJ, Cullen-Unsworth LC, Duarte CM, Eklöf JS, Jarvis JC, Jones BL, Nordlund LM. 2019b. Global challenges for seagrass conservation. Ambio 48: 801–815.

Unsworth RKF, Nordlund LM, Cullen-Unsworth LC. 2019c. Seagrass meadows support global fisheries production. Conservation Letters 12: e12566.

Valdez SR, Zhang YS, Van Der Heide T, Vanderklift MA, Tarquinio F, Orth RJ, Silliman BR. 2020. Positive Ecological Interactions and the Success of Seagrass Restoration. Frontiers in Marine Science 7: 91.

Van Katwijk MM, Thorhaug A, Marbà N, Orth RJ, Duarte CM, Kendrick GA, Althuizen IHJ, Balestri E, Bernard G, Cambridge ML, et al. 2016. Global analysis of seagrass restoration: the importance of largeCscale planting (H Österblom, Ed.). Journal of Applied Ecology 53: 567–578.

Vanstraelen M, Benková E. 2012. Hormonal Interactions in the Regulation of Plant Development. Annual Review of Cell and Developmental Biology 28: 463–487.

Verduin JJ, Seidlitz A, Van Keulen M, Paling EI. 2013. Maximising establishment success of Amphibolis antarctica seedlings. Journal of Experimental Marine Biology and Ecology 449: 57–60.

Waite B, Statton J, Kendrick GA. 2021. Temperature Stratification and Monochromatic Light Break Dormancy and Facilitate On-Demand *In Situ* Germination in the Seagrass *Halophila ovalis*, with Seed Viability Determined by a Novel X-Ray Analysis. Estuaries and Coasts 44: 412–421.

Wang M, Tang X, Zhang H, Zhou B. 2017. Nutrient enrichment outweighs effects of light quality in *Zostera marina* (eelgrass) seed germination. Journal of Experimental Marine Biology and Ecology 490: 23–28.

Waqas M, Korres NE, Khan MD, Nizami A-S, Deeba F, Ali I, Hussain H. 2019. Advances in the Concept and Methods of Seed Priming. In: Hasanuzzaman M, Fotopoulos V, eds. Priming and Pretreatment of Seeds and Seedlings: Implication in Plant Stress Tolerance and Enhancing Productivity in Crop Plants. Singapore: Springer, 11–41.

Waycott M, Duarte CM, Carruthers TJB, Orth RJ, Dennison WC, Olyarnik S, Calladine A, Fourqurean JW, Heck KL, Hughes AR, et al. 2009. Accelerating loss of seagrasses across the globe threatens coastal ecosystems. Proceedings of the National Academy of Sciences 106: 12377–12381.

Xu S, Wang P, Wang F, Zhang X, Song X, Zhou Y. 2021. Responses of eelgrass seed germination and seedling establishment to water depth, sediment type, and burial depth: implications for restoration. Marine Ecology Progress Series 678: 51–61.

Yan A, Chen Z. 2020. The Control of Seed Dormancy and Germination by Temperature, Light and Nitrate. The Botanical Review 86: 39–75.

Yang C-J, Liu Y-S, Liu J, Xu Q, Li W-T, Zhang P-D. 2016. Assessment of the establishment success of *Zostera marina* (eelgrass) from seeds in natural waters: Implications for large-scale restoration. Ecological Engineering 92: 1–9.

Yang YY, Nagatani A, Zhao YJ, Kang BJ, Kendrick RE, Kamiya Y. 1995. Effects of gibberellins on seed germination of phytochrome-deficient mutants of *Arabidopsis thaliana*. Plant & Cell Physiology 36: 1205–1211.

Yang G, Niedziela C, Lu Z. 2005. (176) In Vitro Germination of Galax under Different Culture Medium pH Conditions. HortScience 40: 1052D – 1052.

Zarranz ME, González-Henríquez N, García-Jiménez P, Robaina RR. 2010. Restoration of *Cymodocea nodosa* (Uchria) Ascherson seagrass meadows through seed propagation: seed storage and influences of plant hormones and mineral nutrients on seedling growth in vitro. Botanica Marina 53.

Zhang Y, Xu S, Yue S, Zhang X, Qiao Y, Liu M, Zhou Y. 2022. Reciprocal Field Transplant Experiment and Comparative Transcriptome Analysis Provide Insights Into Differences in Seed Germination Time of Two Populations From Different Geographic Regions of *Zostera marina* L. Frontiers in Plant Science 12: 793060.

Zhu M, Zang Y, Zhang X, Shang S, Xue S, Chen J, Tang X. 2023. Insights into the regulation of energy metabolism during the seed-to-seedling transition in marine angiosperm *Zostera marina* L.: Integrated metabolomic and transcriptomic analysis. Frontiers in Plant Science 14: 1130292.

Ziegler P, Appenroth KJ, Sree KS. 2023. Survival Strategies of Duckweeds, the World’s Smallest Angiosperms. Plants 12: 2215.

